# ATRX safeguards cellular identity during *C. elegans* development

**DOI:** 10.1101/2025.03.11.641662

**Authors:** Janie Olver, Mariya Shtumpf, Karim Hussain, Stephen Methot, Peter Sarkies, Helder Ferreira

## Abstract

ATRX is a member of the SWI/SNF family of ATP-dependent chromatin remodellers. In humans, loss of ATRX function leads to ATRX syndrome, a neurodevelopmental disorder. ATRX mutation in human cell lines is associated with multiple phenotypes including activation of the alternative lengthening of telomere (ALT) pathway, upregulation of retrotransposons and increased sensitivity to replication stress. However, the principal role of ATRX and the reason why its mutation causes such diverse phenotypes is currently unclear. To address this, we studied the role of ATRX in the model organism *Caenorhabditis elegans*. We find that loss of XNP-1, the *C. elegans* homologue of ATRX, recapitulates many human phenotypes. Loss of XNP-1 causes ectopic activation of germline genes in somatic cells, indicating a loss of cellular identity control. Strikingly, mutation of the germline transcription factor *gsox-1* suppresses both this misexpression and multiple *xnp-*1 phenotypes, including developmental delay and telomeric defects. These findings suggest that ectopic germline gene expression underlies the majority of XNP-1-dependent phenotypes, consistent with a role for XNP-1 in maintaining cellular identity, offering insights into the functions of ATRX in humans.

## Introduction

Chromatin structure impacts all processes that require access to DNA. Thus, proteins that alter chromatin structure, particularly ATP-dependent chromatin remodelling enzymes, are important mediators of DNA transcription, replication and repair. Indeed, many disorders are known to be caused by mutation of human ATP-dependent chromatin remodelling genes (1). One of the best described is the neurodevelopmental disorder ATR-X syndrome, which gave its name to the causal gene, the chromatin remodelling factor, ATRX (2).

Fitting with the ubiquitous roles that chromatin remodelling enzymes play in DNA metabolism, ATRX loss is also strongly associated with cancer, specifically those cancers that use the alternative lengthening of telomere (ALT) pathway (3). ALT uses break-induced replication to maintain telomere length (4,5) and ATRX has been shown to act as a tumour suppressor by repressing the ALT pathway (6). Precisely how ATRX influences both ALT and normal development is unclear. This is compounded by the fact that ATRX has been shown to impact many molecular processes. ATRX interacts with DAXX to deposit the replication-independent histone H3.3 at heterochromatin (7,8) and euchromatin (9). Loss of ATRX leads to increased retrotransposon expression (10–12), increased levels of G-quadruplexes (13), reduced sister chromatid cohesion (14,15) and increased sensitivity to replication stress (16,17). Unpicking the relative importance of these molecular functions within a developmental context has been made harder by the fact that complete loss of ATRX is embryonic lethal in standard model organisms such as fruit flies, mice and zebrafish (18–20). Conditional mouse knockouts have allowed the study of ATRX in specific tissues, particularly neuronal lineages (19,21). However, the role of ATRX in early embryogenesis is particularly poorly understood.

To address this, we used the nematode *C. elegans* as a model system, because the loss of its ATRX homolog, XNP-1, does not result in embryonic lethality. *C. elegans* has a particularly well described and stereotypical order of embryonic development where the fate of every cell is known. These decisions are largely governed by the restricted expression of master transcription factors (22,23). However, chromatin structure also plays a complementary role in controlling tissue-specific expression and therefore cellular identity. This is particularly relevant in the very first cell fate decision in *C. elegans* embryogenesis in which germline versus somatic cell fate is specified (24).

In this manuscript, we show that *xnp-1* mutants exhibit ectopic expression of germline genes in somatic cells. We identify the germline transcription factor *F33H1.4* (renamed *gsox-1*) as a suppressor of *xnp-1* sterility at 25°C. Strikingly, *gsox-1* mutation suppresses not only germline misexpression but also seemingly unrelated *xnp-1* phenotypes including developmental delay and telomeric defects. Together, our work identifies a novel role for ATRX homologs in maintaining cellular identity by repressing inappropriate germline gene activation.

## Materials and Methods

### Brood Size

Brood size at 20°C was carried out by singling L4 worms onto OP50 NGM plates and then singling for 3-4 days before counting the total number of progeny. For brood size on plates containing 10 mM hydroxyurea, worms were added to these plates as L1s and then brood size was carried out as above. The worms were maintained on plates containing 10 mM hydroxyurea throughout.

Brood size at 25°C was carried out by shifting L4 worms that were previously grown at 20°C to 25°C. The brood size of their progeny (the F1 or the F2 generation) was recorded in the same way as above.

### C-Circle Assay

Animals were synchronised to L1s and approximately 1000 L1s were seeded onto 5-6 OP50 plates (9 cm). Plates were maintained at 20°C or 25°C for 72 hours until animals reached gravid adult stage.

The C-circle assay was performed as described (25) with the difference that the genomic DNA was extracted by bead-beating. In brief, 300 μl of buffer (100 μg/ml RNase A, 10 mM EDTA, 300 mM NaCl) and 400 μl of 0.5 mm glass beads were added to worm pellets and disrupted by mechanical lysis using a cell homogeniser for 3x 20 s at 6 m/s with cooling on ice in between each step. Proteins were denatured by adding 1% SDS and heating the extracts at 65°C for 10 minutes after which the SDS was precipitated with potassium acetate. The supernatant was collected after spinning and DNA purified using phenol:chloroform:isoamyl alcohol (15:24:1, pH 6.7) extraction and a chloroform back-extraction. DNA was ethanol precipitated and eluted in Tris-EDTA buffer (10 mM Tris-HCl, 100 μM EDTA, pH 7.5). Phi29 polymerase (NEB) was added to 1 μg of genomic DNA in a total volume of 20 µl and incubated at 30°C for 8 hours. This was spotted onto a neutral Hybond-N membrane, 1200J/m2 UV cross-linked, and hybridized with a DIG-labelled (GCCTAA)_4_ probe at 37°C using DIG Easy Hyb (Roche) according to the manufacturer’s instructions.

### Telomere length by terminal restriction fragment (TRF) analysis

Genomic DNA was extracted 1x NTE buffer (100 mM NaCl, 50 mM Tris pH 7.4, 20 mM EDTA), 1% SDS and 500 μg/ml Proteinase K overnight at 65°C. DNA was purified by two consecutive phenol-chloroform washes (15:24:1, pH 6.7), followed by a chloroform back-extraction and ethanol precipitation. DNA was dissolved in Tris-EDTA pH 8 and incubated with 50 μg/ml RNase A for 30 minutes at 37°C followed by a final round of phenol-chloroform extraction and ethanol precipitation. 5 μg purified genomic DNA was digested overnight with HinfI and HaeIII (NEB) at 37°C and resolved on 1% agarose gel in 1x TAE. Following a 20-minute depurination in 250 mM HCl, the gel was washed 2x in denaturing buffer (1.5 M NaCl, 0.5 M NaOH) and 2x in neutralising buffer (1.5 M NaCl, 0.5 M Tris-HCl, pH 8) at room temperature. DNA was transferred onto neutral nylon membrane (Hybond-NX, GE Healthcare) by capillary transfer in 10x SSC buffer (1.5M NaCl, 150 mM sodium citrate, pH 7). After briefly rinsing in 2x SSC buffer (300 mM NaCl, 30 mM sodium citrate, pH 7) DNA was UV crosslinked at 1200 J/m^2^ and hybridised with a digoxygenin-labelled telomere probe (GCCTAA) (1 μg/ml in DIG Easy Hyb^™^ hybridisation buffer, Roche) for 2 hours at 37°C. Membranes were washed 2x in MS buffer (100 mM maleic acid, 150 mM NCl, pH 7.5), 0.3% Tween-20, blocked in MS buffer, 1% BSA, 1% milk, 0.1% Tween-20, probed with AP-conjugated anti-digoxigenin Fab fragments (Roche) at 1:20 000 dilution in the same buffer (1:20000), followed by 3 washes in MS buffer, 0.3% Tween-20. Membranes were equilibrated in AP buffer (100 mM Tris-HCl, pH 9.5, 100 mM NaCl) followed by chemiluminescent detection with CDP-Star^®^ substrate (Roche).

### Quantifying C. elegans growth rates

Post-embryonic growth rates were measured using a luciferase-based assay performed as described previously (26), based on (27). Briefly, wells in luminometer compatible 384- well plates (Berthold Technologies, 32505) were filled with 90 μ S- Basal medium *containing E. coli* OP50 (OD600 = 1) and 100 μM Firefly D-Luciferin (p.j.k., #102111). Single eggs, from animals expressing luciferase under a constitutive and ubiquitous promoter (xeSi312[eft-3p::luc::gfp::unc-54 3′UTR, unc-119(+)] IV) (28), were then loaded into individual wells and plates were sealed with Breathe Easier sealing membrane (Diversified Biotech, BERM-2000). Development was monitored over 96 hrs by measuring luminescence in every well (every 10 mins for 0.5 secs) using a luminometer (Berthold Technologies, Centro XS3 LB 960) placed in a temperature-controlled incubator set to 18.5°C. Luminescence data were analyzed similarly to before (26), but using the Python-based ‘PyLuc’ tool developed by L. Morales Moya (28). The identification of molts (the final phase of each larval stage) was based on a drop in luminescence signal, which results from animals not feeding during this period. For each animal, larval stage durations were then calculated as the time from hatching to the end of the first molt (L1), or from the end of one molt to the end of the subsequent molt (L2 to L4).

### RNA Experiments

RNA extraction was performed as described in (29). Approximately 800 ng of RNA was used as input to each reverse transcription reaction (GoScript Reverse Transcriptase, Promega). The qPCR reaction was carried out on a QuantStudioTM Real-Time PCR system machine (Thermo Scientific) with the following reaction steps: 95°C for 60” ➔ 40x(95°C for 15” ➔ 59°C for 30”). The qPCR primers used are given in Supplementary Table 2.

RNA sequencing of embryos and L1 was carried out at the Oxford Genomics Centre and for the L4 samples was carried out at Novogene, Cambridge. All genotypes were sequenced in biological triplicate. Briefly, reads were trimmed to remove Illumina adaptors using Trim_galore, a wrapper script from Cutadapt (version 0.6.7) (30) using default settings. Trimmed reads were mapped to the reference genome (release WS280, BioProject PRJNA13758) using STAR (version 2.7.10a) (31), without limits on the mapped length and with maximum number of mismatches per pair set to 2. The reads were re-sorted by read name using Samtools (version 1.6) (32). FeatureCounts (Subread, version 2.0.1) (33) was used to generate a count matrix at gene-level. Differential gene expression was conducted in edgeR (version 3.40.0) (34) on a local machine using R (version 4.2.2) applying a minimum gene count of four per replicate and 20 counts across all three replicates per genotype. All scripts can be found in our Github repository (https://gitlab.com/mariya_s/draft_scripts_olver_et_al_2025/-/tree/4aab25604a4a4c716fb24962c6437338bb094249/). SimpleMine web tool from WormBase was used to convert WormBase gene IDs to gene names. Gene set enrichment analysis was also ran on the WormBase web server and the results were plotted using the bubble plot function provided by the SRplot web server (https://www.bioinformatics.com.cn/srplot) (35). Small non-coding RNA sequencing and analysis was performed as described (29). Briefly, 1ug total RNA was treated with RppH (NEB) for 1 hour at 37C to remove 5’ triphosphates. RNA was then extracted by phenol chloroform and precipitated with 3M sodium acetate and ethanol. The resulting RNA was subjected to small RNA sequencing by Oxford Genomics. Fastq files were processed to remove adaptors using Fastx Toolkit. To obtain genome-wide alignments, all reads were aligned to the reference genome (WS280 as above) using Bowtie with the following parameters: -k 1 -v 0 --best to produce alignment sam files, which were further converted to bam files. Bedtools (36) intersect was used to assign reads to different genomic features. 22G-RNAs aligning to transposable elements were specifically interrogated by first selecting reads that were 22 nucleotides long and with a G as the 5’ nucleotide using a custom perl script. These reads were then aligned to the consensus sequences for transposable elements in *C. elegans* downloaded from RepBase (https://www.girinst.org/repbase/) using Bowtie with parameters as above. Antisense reads corresponding to each transposable element were counted using Bedtools genomecov and quantified and visualised using R (version 4.2.2).

### Forward genetic screen and DNA sequencing

Mutagenesis was carried out as described (37). Briefly, synchronized L4 *xnp-1(tm678)* worms were exposed to 50mM ethyl methanesulfonate (EMS) for 4 hours at room temperature, before being washed and placed on OP50 at 20°C until the F1 generation was gravid. At this point ∼250 worms were singled onto new OP50 plates and kept at 25°C. Three fertile lines were isolated and one of these, *xnp-1(hcf3)* was pursued further. *xnp-1(hcf3)* was backcrossed three times to *xnp-1(tm678)* before genomic DNA was extracted (as described in TRF analysis) and sent for sequencing at Novogene (Cambridge Science Park). Potential mutations were mapped, and homozygous mutations identified using the MiModD (version 0.1.9) package developed in the Baumeister lab (https://mimodd.readthedocs.io/en/latest/index.html). The reference genome used was release WS283 (BioProject PRJNA13758).

### Strains

All strains used and their availability are listed in Supplementary Table 1.

## Results

### XNP-1 loss phenocopies several aspects of ATRX loss in human cells

Chromatin remodellers are defined by their ATPase activity and domain swap experiments show that this region dictates remodelling outcomes (38). *C. elegans* XNP-1 and human ATRX share 70% sequence identity within this catalytic domain (Figure 1A), suggesting that despite being half the size of ATRX, XNP-1 likely performs similar chromatin remodelling reactions (39,40).

**Figure One.**
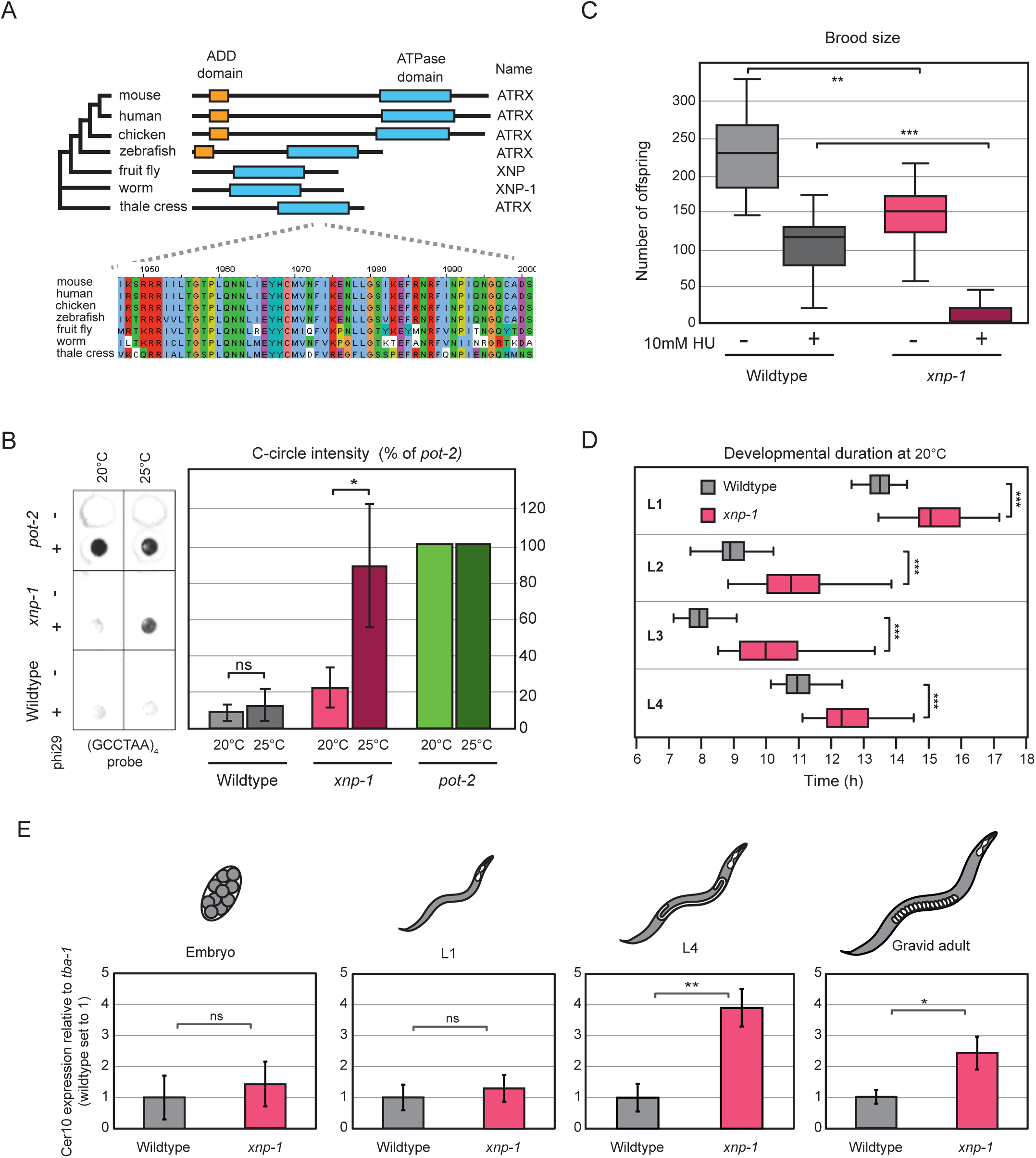
Loss of *C. elegans* XNP-1 mimics several phenotypes of human ATRX mutation. A) XNP-1 is the *C. elegans* homologue of human ATRX. The ADD (ATRX- DNMT3-DNMT3L) domain is not found in ATRX homologues of lower eukaryotes, but the catalytic ATPase domain is highly conserved between humans and worms. (B) *xnp-1* mutant worms display elevated C-circle levels at 25°C. Representative blot from a C-circle assay both at 20°C or 25°C with and without the polymerase Phi29. Quantification of data from five independent C-circle assay experiments using worms synchronised to L1s and then grown for 72 hours at either 20°C or 25°C. Normalised values are presented as percent of *pot-2(tm1400)* signal and the mean across different experiments is shown. Error bars represent the standard deviation from the mean, ns = not significant, * = p < 5 x 10^-2^ (two-tailed Mann-Whitney U test). (C) *xnp-1* mutant worms have lower brood sizes than wildtype and are sensitive to hydroxyurea (HU). Synchronised L1 wildtype or *xnp-1(tm678)* larvae were placed on either OP50 plates or OP50 plates supplemented with 10 mM HU and maintained at 20°C. The brood sizes of 12 - 20 adults per genotype are displayed as box and whisker plots, ** = p < 1 x 10^-3^, *** = p < 1 x 10^-5^ (Welch’s test). (D) *xnp-1* mutant worms develop slower than wildtype. Synchronised L1 wildtype or *xnp-1(tm678)* larvae were singled into individual wells and imaged over two days to monitor growth. Box and whisker plots of data from at least 70 worms per genotype indicating the total time to complete each larval stage, *** = p < 1 x 10^-6^ (two tailed T-test). (E) The LTR retrotransposon *Cer10* is de-repressed most strongly in larval stages of *xnp-1* animals that contain a germline. RT-qPCR of *Cer10* with *tba-1* used as the reference in wildtype and *xnp-1(tm678)* animals with *Cer10* expression normalised to wildtype. The mean and standard deviation is shown for three biological replicates. ns = not significant, * = p ≤ 5 x 10^-2^, ** = p ≤ 10 x 10^-3^ (two tailed T-test).

To study the effects of XNP-1 loss, we used a null mutant, *xnp-1(tm678)*. This allele contains a 673bp deletion in the *xnp-1* gene and additionally introduces a premature stop codon before the ATPase domain (39,40). Loss of ATRX in human cells has been linked to several phenotypes such as ALT activation, sensitivity to replication inhibitors, development delay and de-repression of retrotransposons (3,12,16). We therefore tested whether loss of XNP-1 induced similar phenotypes in *C. elegans*.

Human ALT positive tumours are defined clinically by the presence of C-circles: partially single-stranded circles of extrachromosomal telomeric DNA (25). The connection between ALT and C-circles extends beyond humans as *C. elegans* ALT-like strains also show increased C-circle levels (41–43). When we examined synchronized populations of wildtype and *xnp-1* gravid adults grown at 20°C, we detected no significant differences in C-circle levels. However, when grown at 25°C, a known stress condition, *xnp-1* showed significantly elevated C-circle levels compared to wildtype (Figure 1B). These levels were comparable to those in *pot-2(tm1400)* mutants, which form long-term ALT survivors in telomerase-null backgrounds (42,43). This stress-dependent C-circle induction parallels the human situation, where additional factors such as telomeric DNA breaks (44) or protein-DNA adducts (45) are required for robust C-circle formation in ATRX mutant cells.

ATRX mutant cells show signs of replication stress and are sensitive to hydroxyurea (HU) (16). This drug induces replication stress by reducing the levels of deoxyribonucleotides in S phase, thereby uncoupling the movement of DNA polymerases from the replicative CMG helicase. We exposed worms to 10mM hydroxyurea from the L1 stage onwards and quantified its effect by measuring brood size of gravid adults two days later. Even without HU treatment, *xnp-1* worms had significantly smaller brood sizes than wildtype (Figure 1C). However, *xnp-1* mutants showed markedly increased HU sensitivity compared to wildtype. While 10mM HU caused an approximately two-fold decrease in wildtype brood size, it resulted in a more than nine-fold drop in *xnp-1*, rendering them nearly sterile (Figure 1C).

ATRX syndrome causes neurodevelopmental delay in humans (46). As this is difficult to assay in worms, we instead assayed whether XNP-1 affects the rate of post-embryonic development. We observed that *xnp-1* mutants took significantly longer than wildtype to develop at every larval stage (Figure 1D). Moreover, the normally tight synchrony of worm development was weakened in *xnp-1* (Supplementary Figure S1).

Loss of ATRX leads to increased expression of endogenous retroviruses (ERVs) in mammalian stem cells (12). To test whether XNP-1 similarly regulates transposable elements, we examined both individual retrotransposons and genome-wide transposon expression. We focused on Cer10, an LTR retrotransposon for which validated qPCR primers are available (47), as LTR retrotransposons are the elements in worms most similar to mammalian ERVs (48). When we examined Cer10 expression across developmental stages, levels were similar between *xnp-1* and wildtype in embryos and L1 larvae but were significantly elevated in *xnp-1* L4s and adults (Figure 1E). RNA-seq analysis revealed that retrotransposon upregulation was not restricted to later stages. Multiple transposon families were significantly upregulated in *xnp-1* mutant embryos and L1 larvae, including other LTR retrotransposons and PIF-Harbinger DNA transposons (Supplementary Figure S2A). Small RNA sequencing revealed no differences in 22G-siRNA levels targeting these transposons, indicating that XNP-1 regulates retrotransposons independently of the siRNA pathway (Supplementary Figure S2B).

To determine whether XNP-1 silences retrotransposons through H3K9me3-dependent mechanisms, we examined the role of SET-25, the only *C. elegans* H3K9 trimethylase (49). Consistent with previous data (56), *set-25* single mutants showed increased Cer10 retrotransposon expression. However, *xnp-1; set-25* double mutants had significantly higher Cer10 levels than *set-25* single mutants (Supplementary Figure S3), indicating that XNP-1 and SET-25 function in independent pathways.

Overall, our data demonstrates that loss of XNP-1 in *C. elegans* recapitulates multiple key phenotypes seen in ATRX mutant human cells, supporting functional conservation despite structural differences between organisms.

### XNP-1 represses germline genes distinct from SynMuvB

Analysis of polyA-enriched RNA sequencing data revealed relatively few differentially expressed genes in either *xnp-1* embryos or L1 larvae when grown at 20°C (Figure 2A and Supplementary Table 3). Downregulated genes between *xnp-1* embryo and L1 larvae were largely distinct (Figure 2B) but tissue enrichment analysis displayed an association with the intestine (Supplementary figure S4A). In contrast, there was a large degree of overlap between genes upregulated in embryos and L1s (Figure 2B). Tissue enrichment analysis revealed that these genes were very strongly associated with the germline (Figure 2C). This was surprising as neither embryos nor starved L1 larvae have a germline *per se*. Indeed, the only two germline progenitor cells (Z2 and Z3) in both these developmental stages are maintained in a transcriptionally quiescent state (50). Therefore, the increased level of germline expression we see in *xnp-1* embryos and L1 larvae is most likely to be due to ectopic expression of germline genes from somatic cells. The ectopic activation of germline genes in somatic cells indicates a loss of cellular identity control. This suggests XNP-1 normally functions to maintain the transcriptional boundary between germline and somatic cell fates by repressing germline-specific genes in somatic cells. This resembles the phenotype caused by loss of SynMuvB genes (51), sometimes referred to as the DREAM complex in other species (52). A hallmark of synMuvB mutation in worms is increased expression of PGL-1, a P granule component found in all germ cells (51). We monitored *pgl-1* expression in *xnp-1* embryos or L1 larvae at 25°C, a condition in which the increase in PGL-1 in SynMuvB mutants is particularly pronounced (51). However, we found no significant increase in *pgl-1* mRNA levels in *xnp-1* compared to wildtype control (Figure 2D), suggesting that *xnp-1* may not be a SynMuvB gene. Indeed, when we looked beyond *pgl-1* to include all genes that become upregulated in *lin-35,* a key member of the SynMuvB family (51), we saw very little overlap with genes that become upregulated in *xnp-1* (Supplementary Figure S4B). Another hallmark of SynMuvB mutants is that they display larval arrest at elevated temperatures, which is suppressed by RNAi of the H3K36 methyltransferase *mes-4* (51). However, loss of XNP-1 did not show a larval arrest phenotype at 25°C (Figure 2E) and *mes-4(RNAi)* did not rescue the reduced brood size *xnp-1* at 25°C (Figure 2F and Supplementary Figure S4C). Altogether, these data show that XNP-1 and SynMuvB repress germline genes through distinct pathways. Alongside the upregulation of germline genes, we observed downregulation of somatic genes in *xnp-1* embryos and L1 larvae.

**Figure Two.**
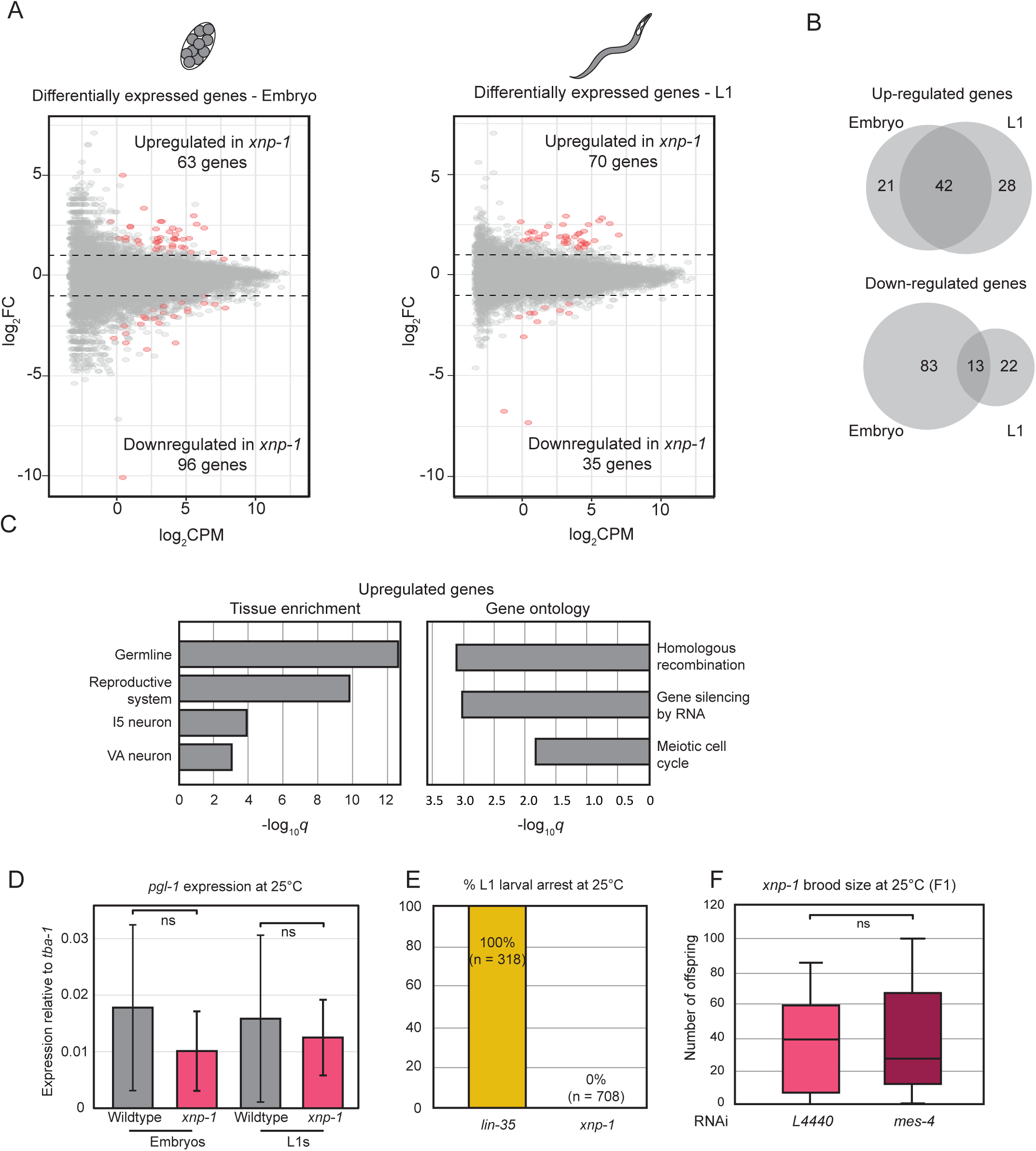
XNP-1 prevents ectopic germline expression in a different manner to the SynMuvB pathway. (A) Loss of XNP-1 leads to changes to a modest number of differentially expressed genes (DEGs) in both embryos and L1s grown at 20°C. Genes whose expression is significantly altered in *xnp-1* are marked in red within the MA plot, false discovery rate (FDR) < 0.05. (B) Upregulated genes in *xnp-1* are more likely to behave similarly in both embryo and L1 samples compared to downregulated genes. (C) Analysis of genes that are significantly upregulated in *xnp-1* samples (from part A) indicates that they are normally expressed within the germline and typically have functions in DNA repair and meiosis. (D) *pgl-1* is not upregulated in *xnp-1* mutants. RT-qPCR of *pgl-1* normalised to *tba-1* in wildtype and *xnp-1(tm678)* embryos and L1 larvae grown at 25°C. The mean and standard deviation is shown for three biological replicates. ns = p > 5 x 10^-2^ (two tailed T-test). (E) *xnp-1* animals (in contrast to *lin-35*) do not show L1 larval arrest at 25°C. Synchronised L4s were placed at 25°C and left to produce offspring. The number of offspring scored for an arrest phenotype and the percentage of L1 larval arrest is indicated. (F) The low brood size of *xnp-1* animals at 25°C is not rescued by *mes-4(RNAi).* Synchronised *xnp-1* L1s were placed at 25°C onto control or *mes-4(RNAi)* plates. Once they developed to gravid adults, they were singled and the brood size of those adults (F1) scored. Box and whisker plots from 15 - 20 adults are displayed, ns = not significant.

### Identification of gsox-1 as a suppressor of xnp-1 sterility at 25°C

To identify genes that function with XNP-1, we turned to genetic suppressor analysis. Forward genetic screens reveal functionally relevant genes in living animals, where transcription, development, and stress responses are tightly integrated. *C. elegans* lacking XNP-1 become completely sterile within two generations when grown at 25°C (Figure 3A), providing a robust phenotype for screening. We used ethyl methanesulfonate (EMS) to mutagenise *xnp-1* mutants and screen for suppressor mutants that restored fertility at 25°C (Figure 3B). One suppressor, *xnp-1(hcf3)*, could be maintained indefinitely at 25°C with significantly larger brood sizes than the original *xnp-1(tm678)* strain (Figure 3C). Genome sequencing of this strain identified a locus on chromosome II containing G/C to A/T mutations characteristic of EMS mutagenesis. Within this locus were four genes (*toe-1*, *mog-5*, *hgap-2* and *F33H1.4*) with single point mutations within their exons that resulted in protein-coding changes. To definitively link the suppressor phenotype to a single mutation, we independently introduced each of these mutations using CRISPR into a wildtype background and then crossed them into unmutagenised *xnp-1(tm678).* Only the *F33H1.4* mutation suppressed *xnp-1* sterility at 25°C (Figure 3D). This gene encodes a nematode-specific transcription factor expressed in the germline that binds to germline gene promoters (53,54). Based on this, we re-named *F33H1.4* as *gsox-1* (germline suppressor of *xnp-1*).

**Figure Three.**
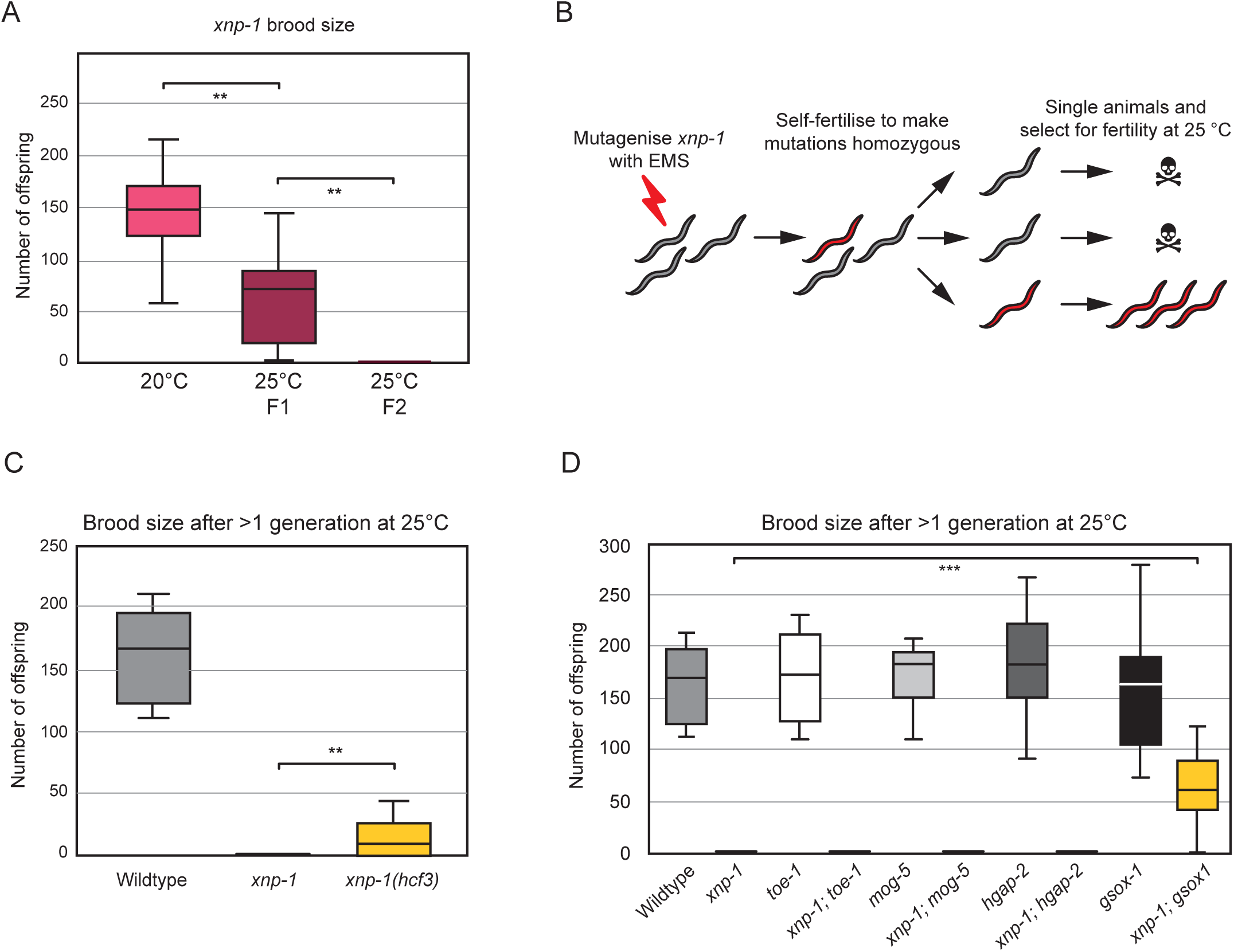
Identification of *gsox-1* as a suppressor of *xnp-1* sterility at 25°C. (A) Loss of XNP-1 leads to rapid sterility at 25°C. Synchronised *xnp-1* L1s were placed at 25°C onto OP50 plates. Once they developed to gravid adults, they were singled and the brood size of those adults (F1) scored. The subsequent generation at 25°C (F2) were sterile. In contrast, *xnp-1* worms can be maintained indefinitely at 20°C. Box and whisker plots from 15 - 20 adults are displayed, ** = p ≤ 10 x 10^-4^ (two tailed T-test). (B) Schematic of forward genetic screen to isolate *xnp-1* suppressors. (C) Identification of *xnp-1(hcf3)* as a weak suppressor of temperature induced sterility. Box and whisker plot of the number of F2 progeny of wildtype, the original *xnp-1(tm678)* strain and the suppressor line *xnp-1(hcf3)*, * = p < 5 x 10^-2^ (Welch’s test), n > 19 for all genotypes. (D) *gsox-1* is the causal gene behind the fertility of *xnp-1(hcf3)* at 25°C. Box and whisker plot of the number of F2 progeny of wildtype, *xnp-1(tm678)*, *toe-1(syb6057)*, *mog-5(syb5976)*, *hgap-2(syb7188)*, *gsox-1(syb7245)* and double mutants of these with *xnp-1(tm678)*, *** = p < 1 x 10^-4^ (Welch’s test), n > 10 for all genotypes.

### Mutation of *gsox-1* suppresses the majority of misregulated germline expression in *xnp-1*

The connection of *gsox-1* to germline gene expression was intriguing given the ectopic germline gene expression seen in *xnp-1* mutants at 20°C (Figure 2A-C). We therefore examined transcriptional changes in *xnp-1* animals at 25°C, the condition causing rapid sterility. Wildtype and *xnp-1* mutant embryos were hatched at 20°C and then grown from L1 to L4 larvae at 25°C (Figure 4A). Transcriptomic analysis revealed that >6000 genes were significantly altered (FDR <0.05) compared to wildtype (Figure 4A), far more than observed at 20°C (Figure 2A).

**Figure Four.**
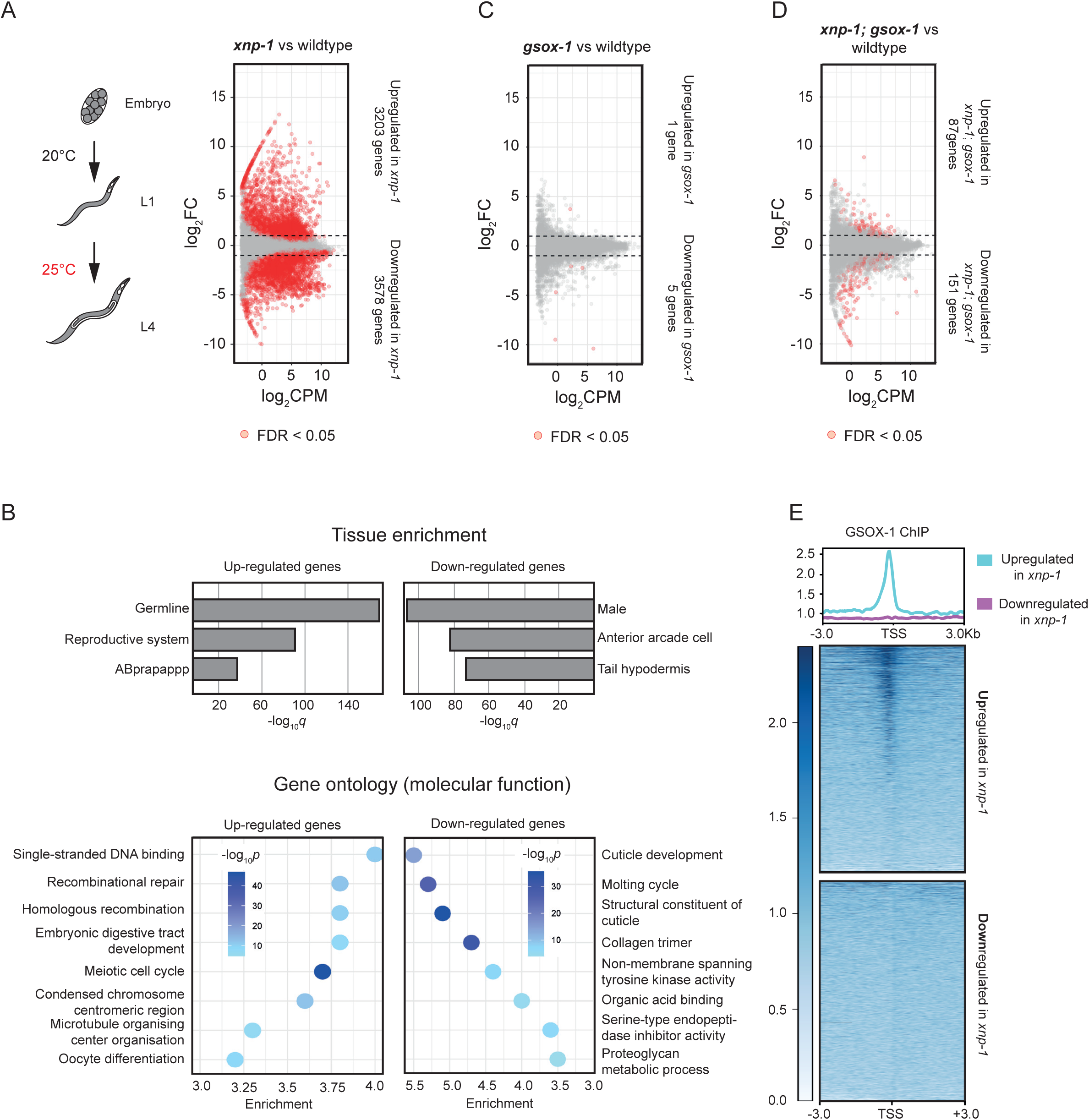
Mutation of *gsox-1* suppresses the majority of transcriptional changes in *xnp-1*. (A) Schematic of growth conditions of wildtype and *xnp-1* worms prior to processing for polyA-RNA sequencing. MA plot illustrating that a very large number of transcripts are misrergulated in *xnp-1* relative to wildtype. Differentially expressed genes (DEGs) were defined using a false discovery rate (FDR) of less than 0.05. (B) Upregulated and downregulated DEGs are associated with distinct tissues and molecular functions. Upregulated DEGs in *xnp-1* are typically expressed in the germline (similar to what is seen in Figure 2C). In contrast, downregulated DEGs are typically expressed in males and the hypodermis. Enriched gene ontology terms for up- and downregulated DEGs are ordered by enrichment over the background set (all genes detected) and colour coded by their FDR. Upregulated DEGs in *xnp-1* are typically associated with functions in DNA repair and meiotic cell cycle (again similar to what is seen in Figure 2C). In contrast, downregulated DEGs have functions in cuticle formation and moulting. (C) Mutation of *gsox-1* by itself does not cause significant transcriptional disruption. (D) Mutation of *gsox-1* reduces the number of DEGs in an *xnp-1* background. MA plot, as described in part A, showing that the number of DEGs in *xnp-1; gsox-1* is more than 25-fold lower than in *xnp-1*. (E) GSOX-1 shows robust binding at promoters of genes upregulated in *xnp-1* but not at downregulated genes. ChIP-seq data from (53) analysed using deepTools. Top panel: Average GSOX-1 signal across all genes in each category, cantered on the TSS +/- 3kb. Bottom panel: Heatmap showing GSOX-1 binding at individual genes, with each row representing one gene.

More importantly, when we looked at the identity of upregulated genes, we saw that they were again very strongly associated with the germline and the meiotic cell cycle (Figure 4B), similar to what was seen at 20°C (Figure 2C). This meant that the same types of genes were affected in *xnp-1* worms at both 20°C (relatively normal physiology) and 25°C (profound developmental defects), but that the scale of the transcriptional changes was much larger at 25°C. Altogether, this suggested a link between germline upregulation and sterility in *xnp-1*. Down-regulated genes in *xnp-1* were associated with skin (hypodermis), the cuticle and moulting cycle (Figure 4B). Cuticle formation and the moulting cycle drive the growth of larval stages in *C. elegans*. It is possible that the disruption of these pathways contributes to the slow post-embryonic development of *xnp-1* animals (Figure 1D).

We observed very few significant transcriptional changes in our single *gsox-1* mutant compared to wildtype (Figure 4C). This is consistent with the *gsox-1* allele (R1192Q) being a hypomorph since RNAi knockdown of *gsox-*1 is embryonic lethal (55). Despite having a minimal effect in wildtype, *gsox-1* mutation had a marked effect in an *xnp-1* background, supressing the vast majority of transcriptional misregulation. The double *xnp-1; gsox-1* mutant had only 234 differentially expressed genes (DEGs) relative to wildtype (Figure 4D), which is much less than the 6781 DEGs between *xnp-1* and wildtype (Figure 4A).

To distinguish between direct and indirect effects, we leveraged existing ChIP-seq data for GSOX-1 in wildtype young adults (53). GSOX-1 displayed robust binding at the promoters of upregulated genes but essentially no binding at *xnp-1* downregulated genes (Figure 4E). This was specific to GSOX-1 as we did not see the same binding pattern in the unrelated transcription factor NHR-23 (Supplementary figure S5). This enrichment is consistent with GSOX-1’s known role as a germline transcription factor (54) and suggest a hierarchy of effects: *gsox-1* directly suppresses *xnp-1* upregulated (germline) genes, whereas its effects on downregulated genes are indirect. This argues that ectopic germline gene expression is the primary defect in *xnp-1*, with disruption of somatic programs as downstream consequences and is consistent with the pattern of misregulation seen in embryos and L1 larvae at 20°C. Accordingly, these data support a direct link between fertility defects in *xnp-1* and germline gene misregulation.

### Altered cellular identity likely drives the developmental and telomeric phenotypes of *xnp-1*

We wondered whether *gsox-1* could also suppress other *xnp-1* phenotypes. Indeed, we found that *gsox-1* significantly suppressed the slow growth of *xnp-1* at 20°C. The *xnp-1; gsox-1* double mutant developed significantly faster than *xnp-1* at every larval stage (Figure 5A). This included the first larval stage, L1, which has no significant germline and only two germline progenitor cells per animal. Therefore, *gsox-1* can suppress *xnp-1* developmental phenotypes that are not directly linked to the germline. This interpretation was strengthened when we looked at telomeric phenotypes. We observed that *gsox-1* partially suppressed the long telomere phenotype of *xnp-1* animals (Figure 5B) and almost completely supressed the increased C-circle levels at 25°C (Figure 5C). Similarly, *gsox-1* largely suppressed the upregulation of the Cer10 retrotransposon in *xnp-1* L4 larvae at 25°C (Figure 5D). Our data show that these seemingly unconnected *xnp-1* mutant phenotypes are all rescued by mutation of a germline transcription factor. This raises the possibility that these *xnp-1* phenotypes are driven by altered cellular identity, specifically an increase in a germline-like cell fate.

**Figure Five.**
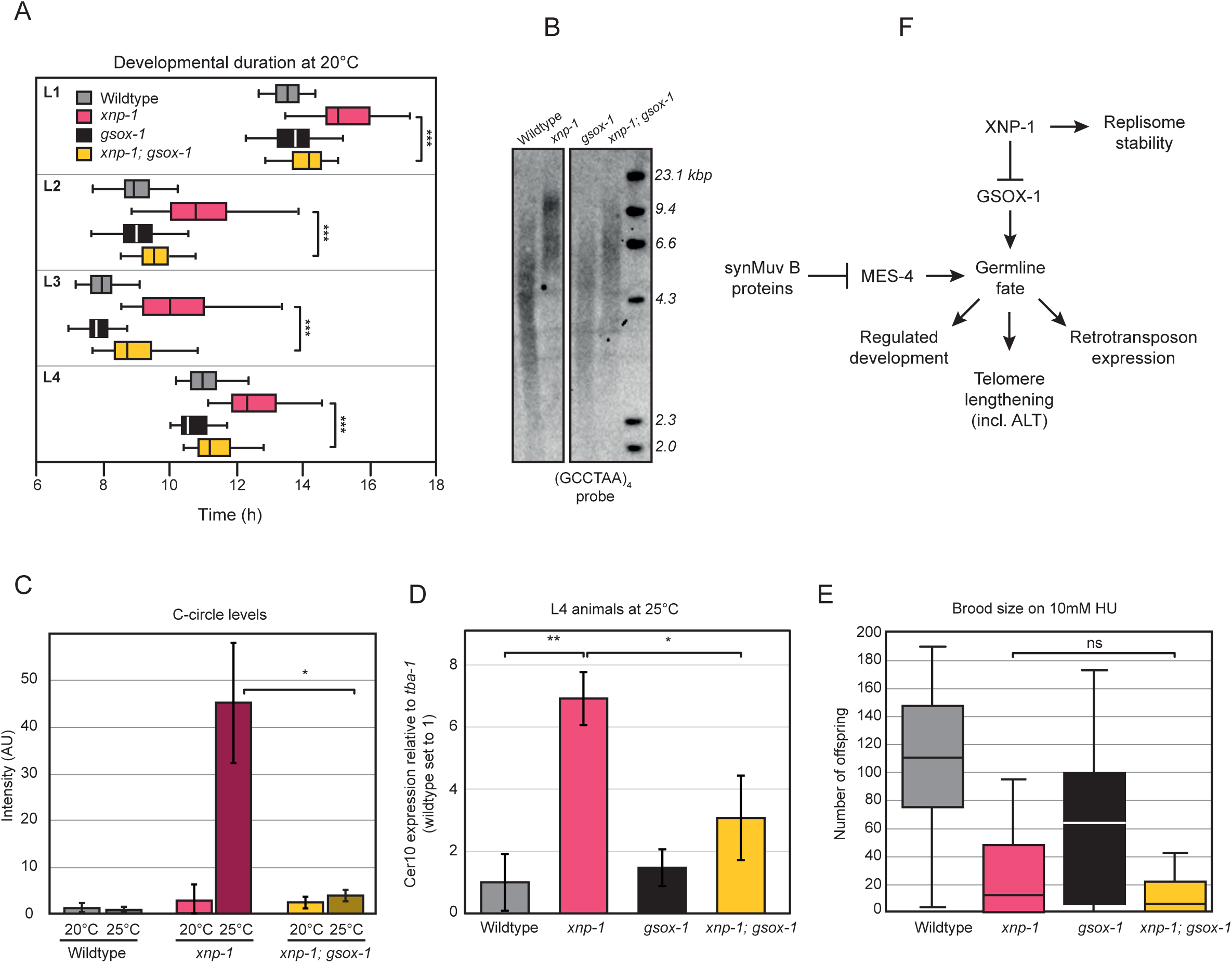
The developmental and telomeric phenotypes of *xnp-1* are linked and likely due to altered cellular identity. (A) *gsox-1* partially recues the slow development phenotype of *xnp-1* at 20°C. Synchronised L1 larvae were singled into individual wells and imaged over two days to monitor growth. Data for wildtype and *xnp-1* as in Figure 1D. Box and whisker plots of data from at least 70 worms per genotype indicating the total time to complete each larval stage, *** = p < 1 x 10^-6^ (two tailed T-test). (B) *xnp-1* worms have longer telomeres than wildtype which is partially suppressed by *gsox-1*. Southern blot, probed with a telomere-specific sequence, of an asynchronous population of the indicated strains grown at 20°C. (C) *gsox-1* recues the elevated C-circle levels of *xnp-1* at 25°C. Quantification of data from five independent C-circle assay experiments using worms synchronised to L1s and then grown for 72 hours at either 20°C or 25°C. Data for wildtype and *xnp-1* as in Figure 1B, * = p < 5 x 10^-2^ (two-tailed Mann-Whitney U test) (D) *gsox-1* partially recues the increased *Cer10* expression observed in *xnp-1.* RT-qPCR of *Cer10* levels normalised to *tba-1* in L4 stage animals of the indicated genotypes grown at 25°C from L1 onwards. The level of wildtype Cer10 expression is set to 1. The graph displays the mean and standard deviation from three biological replicates. * = p ≤ 5 x 10^-2^, ** = p ≤ 5 x 10^-3^ (two tailed T-test). (E) *gsox-1* does not supress the HU sensitivity of *xnp-1.* Synchronised L1 larvae of the indicated genotypes were placed on either OP50 plates or OP50 plates supplemented with 10 mM HU and maintained at 20°C. Data for wildtype and *xnp-1* as in Figure 1C. The brood sizes of 12 - 20 adults per genotype are displayed as box and whisker plots, ns = not significant. (F) Model for how XNP-1 works in *C. elegans*. XNP-1 represses the ectopic expression of germline genes in a manner distinct form the SynMuvB pathway. The inappropriate activation of a germline-like fate is the likely cause of the developmental, telomeric and retrotransposon repression defects seen in in *xnp-1* as all these phenotypes are suppressed by *gsox-1*.

The one *xnp-*1 phenotype which *gsox-1* could not suppress was sensitivity to HU (Figure 5E). We observed that the brood size of *xnp-1; gsox-1* double mutants on 10mM HU was no higher than *xnp-1*. Instead, *gsox-1* mutants were themselves mildly sensitive to HU. Thus, XNP-1’s role in replisome stability operates independently of GSOX-1 and is unlikely to be linked to germline identity, unlike its other functions described here (Figure 5F).

## Discussion

The ATRX gene has long been linked to human disease; it is the causal mutation behind ATRX syndrome (2) and is mutated in 90% of ALT positive cancers (3). In this study, we find that loss of the ATRX homologue, XNP-1 in *C. elegans*, leads to many of the same phenotypes in nematodes as in human cells, including: de-repression of the ALT pathway, increased sensitivity to replication stress, de-repressed retrotransposon expression and disrupted development. Given that nematodes and humans diverged over a billion years ago (56), this suggests that ATRX syndrome and upregulation of ALT are caused by disruption of an evolutionarily ancient function of the ATRX gene.

One of the striking aspects of ATRX has been the multiple, seemingly disparate, phenotypes observed when it is mutated (57). It has not been clear how these various ATRX mutant phenotypes are linked, or whether they instead represent separate, independent functions. The fact that we see conservation of so many ATRX phenotypes in *C. elegans*, hints at a common underlying mechanism. This is strengthened by the fact that a single suppressor mutant of *xnp-1* sterility (*gsox-1)* also supressed other *xnp-1* phenotypes that it was not selected for. The simplest explanation for this observation is that these different *xnp-1* phenotypes are all distinct downstream consequences of a common upstream problem.

Given that GSOX-1 is a germline transcription factor (53,54), we propose that the initial trigger in *xnp-1* mutants is misregulated cellular identity, more specifically due to cells taking on a more germline-like fate. Although GSOX-1 functions primarily in the germline, it is also expressed in some somatic tissues, such as glia (58). Thus, reducing GSOX-1 levels likely suppresses inappropriate germline programs in both tissues. GSOX-1 binds robustly at the promoters of genes that are upregulated in *xnp-1* mutants (germline genes) but shows no binding at downregulated genes (somatic genes) in wildtype. This indicates that GSOX-1 directly activates these germline genes under normal conditions. Consequently, when *gsox-1* is mutated in an *xnp-1* background, the suppression of these upregulated genes restores proper germline identity, with down-regulation of soma-specific genes being indirect secondary consequences.

How does loss of cellular identity control explain the diverse *xnp-1* phenotypes we observe? Disruption of gene regulation within the germline is likely to impair its development, which can directly reduce fertility. Ectopic expression of germline transcripts in somatic cells could also explain the slower development of *xnp-1* animals during developmental stages (L1) where they do not contain a germline *per se* (Figure 1D). Since germline genes are upregulated in *xnp-1* (Figure 2A and 4A), and retrotransposons normally increase during meiosis (59,60), the Cer10 upregulation we observe is consistent with an ectopic meiotic environment caused by inappropriate germline gene expression. Similarly, amongst the genes whose expression increases significantly in *xnp-1* L4 larvae at 25°C are the telomerase catalytic subunit (*trt-1*), the telomere shelterin components (*pot-1* and *pot-2)* as well as several DNA repair proteins (Supplementary Table 3). Since ALT is driven by activation of inappropriate DNA repair pathways at dysfunctional telomeres (5), aberrant germline gene expression could pre-dispose telomeres to inappropriate lengthening and dysfunction. However, not all *xnp-1* phenotypes are explained by loss of cellular identity control. We find that *gsox-1* does not suppress the HU sensitivity of *xnp-1* (Figure 5E). This suggests that the fork stability phenotype of *xnp-1* represents a functionally distinct role of XNP-1 (Figure 5F).

Most models of ALT invoke aberrant DNA repair at a damaged replication fork (61). However, our data indicates that the fork stability phenotype of *xnp-1* (HU sensitivity) is not linked to other *xnp-1* phenotypes. At first glance this is surprising as expressing meiotic proteins in human cells is sufficient to induce replication stress (62). It is possible that XNP-1 plays a more direct role at the replication fork. Human ATRX associates physically with the MCM complex (63), perhaps pointing to a mechanism by which it travels with the fork to promote replisome stability.

The precise mechanism by which XNP-1 controls gene expression is not clear. Our data on retrotransposon expression indicated that XNP-1’s effect does not function via the siRNA pathway and is independent of H3K9me3-dependent silencing. As a member of the ATP-dependent family of chromatin remodeling enzymes, ATRX may play a role in regulating the chromatin template. However, interestingly, ATRX is also involved in functions beyond chromatin formation (64) and a recent study found that most genes whose expression increased following ATRX mutation did not show a corresponding increase in promoter chromatin accessibility (65). This suggests that ATRX may not regulate transcription merely via heterochromatin formation.

Our findings in *C. elegans* are consistent with observations in mammalian systems. ATRX is involved in regulating gene expression and maintaining cellular identity in mammalian cells. Loss of ATRX leads to altered differentiation of mouse oligodendrocyte and human mesenchymal progenitor cells (65,66). Moreover, a mouse model of ATRX syndrome shows signs of early developmental defects (67). Thus, loss of ATRX leads to changes in stem cell identity and a reduction in their ability to resist differentiation into inappropriate cell fates. This connection might also be relevant to ATRX’s role in supressing ALT. A recent study shows that induction of differentiation is a key trigger for ALT in ATRX-deficient stem cells (68). ALT is more frequent in childhood cancers (69). In these cancers, cells have less time to accumulate additional cancer-driving mutations, suggesting that changes in their initial cellular identity may play a more significant role in their eventual malignant transformation.

Mutation of *xnp-1* manifests as a large increase in germline gene expression (Figure 3C, 5B). However, this may be because germline is hypothesized to be the default cell fate in *C. elegans* (70). Thus, XNP-1 may be required to restrict cell fate specification more broadly rather than specifically repress germline genes *per se*. In *C. elegans*, GSOX-1 serves as the key downstream effector of this identity control, yet this transcription factor is not conserved beyond nematodes, whereas XNP-1/ATRX is. This suggests that while the specific transcriptional effectors may differ across species, preventing inappropriate cell fate specification is a fundamental and conserved role of ATRX.

## Acknowledgements

Some strains were provided by the Caenorhabditis Genetics Center, which is funded by the National Institutes of Health Office of Research Infrastructure Programs (P40 OD010440). We thank Prof. Julie Ahringer and Dr Chantal Wicky for helpful discussions. The authors acknowledge Research Computing at the James Hutton Institute for providing computational resources and technical support for the UK’s Crop Diversity Bioinformatics HPC (BBSRC grants BB/S019669/1 and BB/X019683/1), use of which has contributed to the results reported within this paper. MS was supported by an EASTBIO studentship, (BBSRC) [grant number BB/M010996/1]. We thank Peter Thorpe and Marco Ferreira Fernandes for help with bioinformatics.

