## Supplementary figures and images for "ATRX safeguards cellular identity during *C. elegans* development"

### supplementary figures

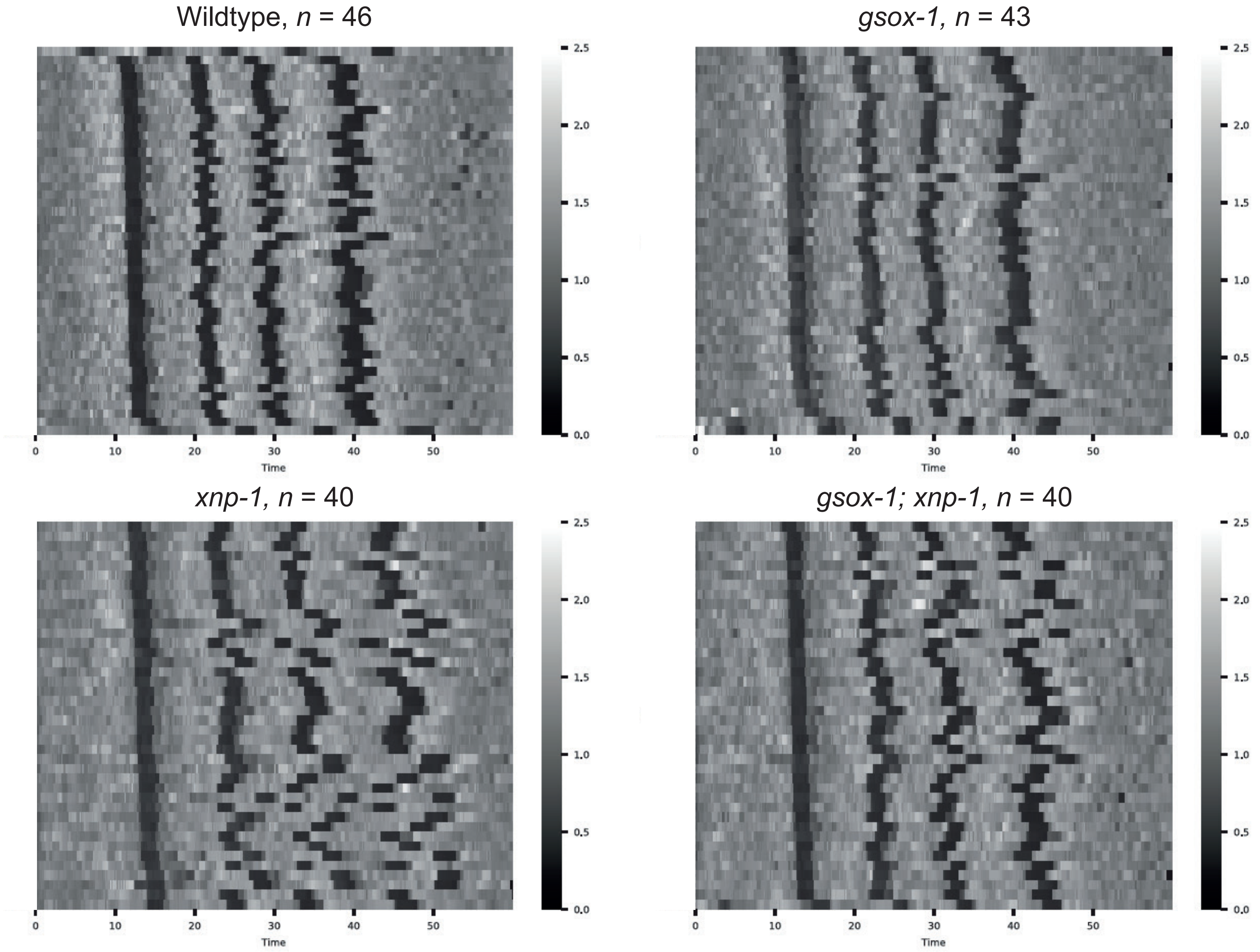

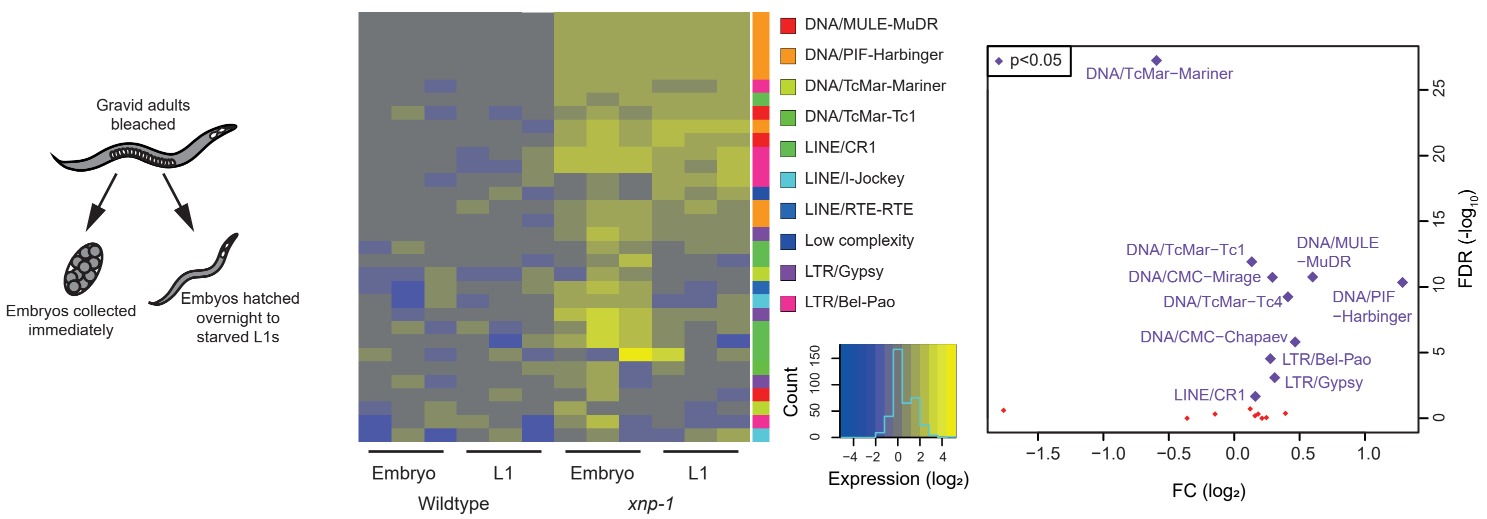

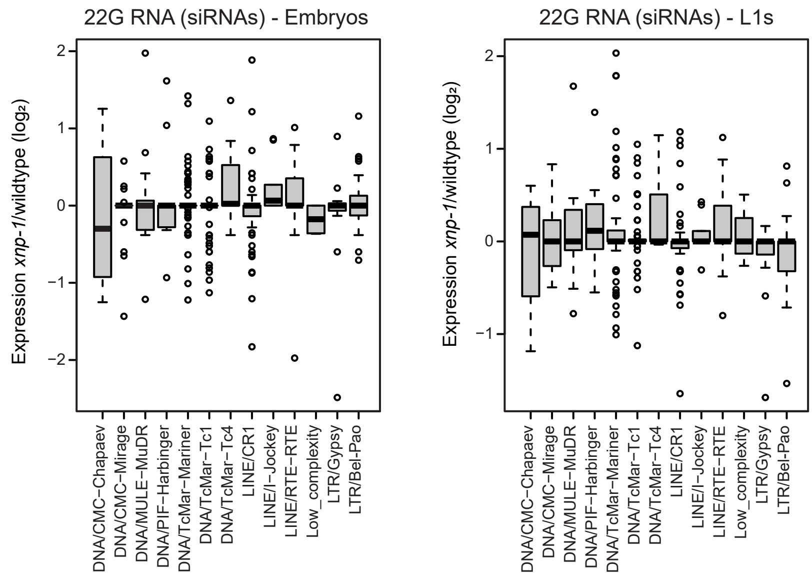

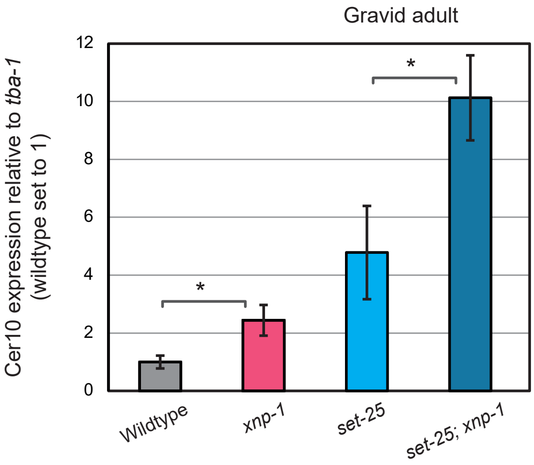

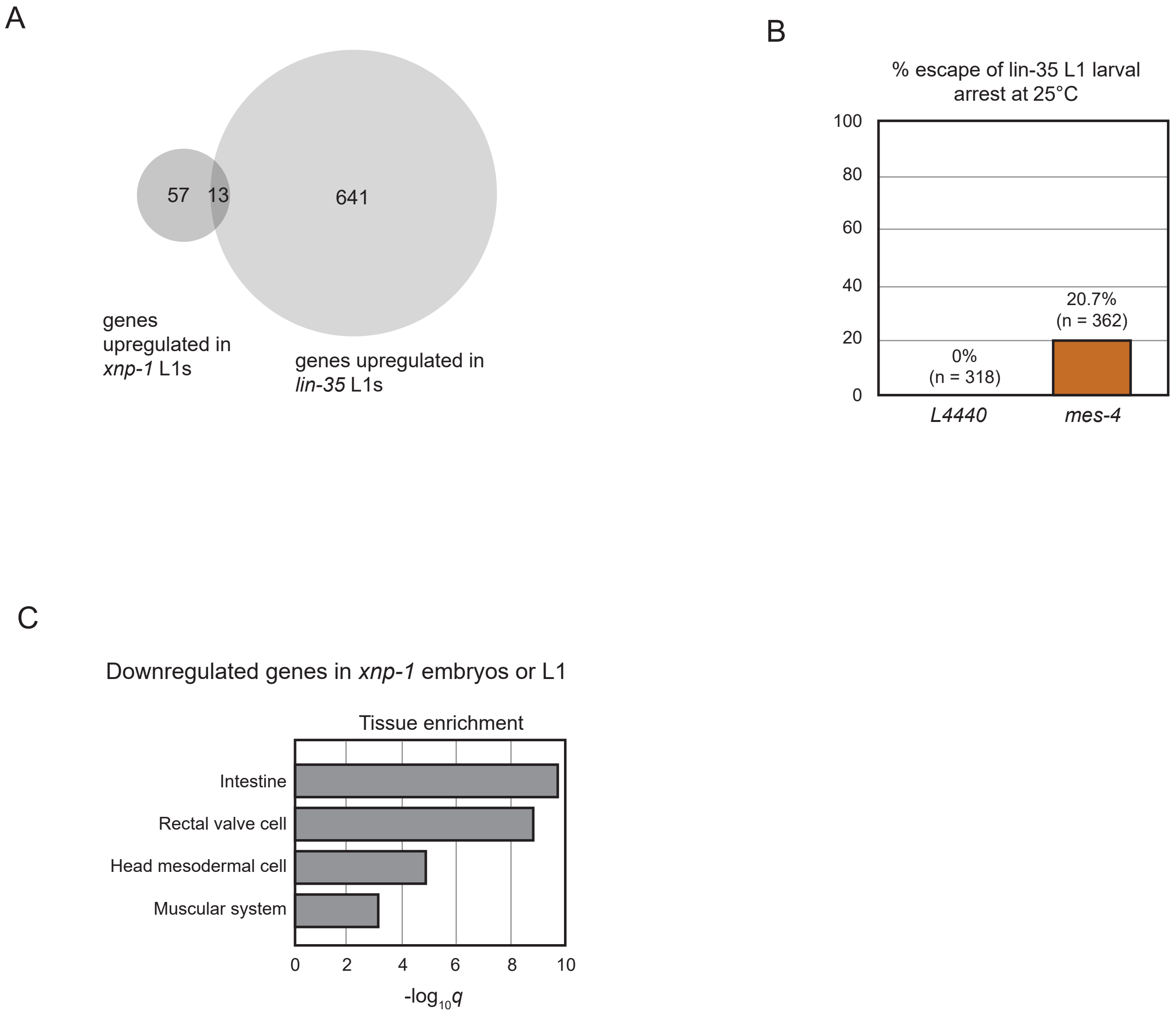

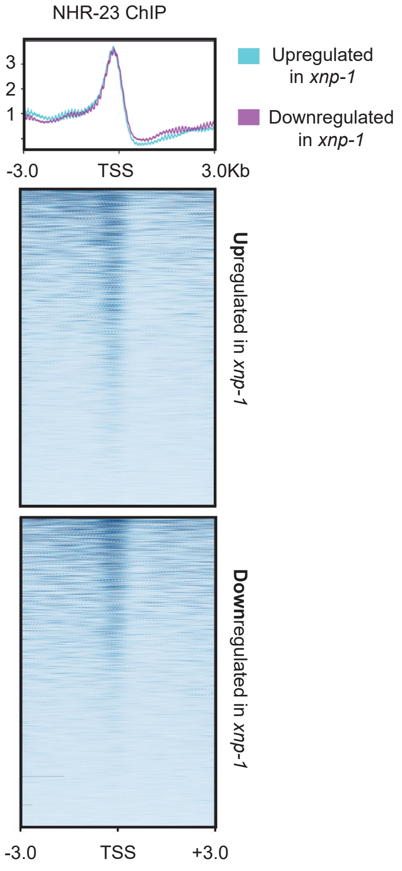
