## supplementary table 1 for "ATRX safeguards cellular identity during *C. elegans* development"

**Supplementary Table One**

|  | **Strain Name** | **Genotype** | **Source** |
| --- | --- | --- | --- |
| Wildtype | N2 | *Bristol strain N2* |  |
| *xnp-1* | HFW4 | *xnp-1(tm678)* | CGC |
| *pot-2* | HFW2 | *pot-2(tm1400)* | CGC |
| *set-25* | HFW62 | *set-25(n5021)* | CGC |
| *xnp-1; set-25* | HFW111 | *xnp-1(tm678); set-25(n5021)* | This study |
| *his-72* | HFW91 | *his-72(tm2066)* | CGC |
| *his-72; xnp-1* | HFW137 | *his-72(tm2066); xnp-1(tm678)* | This study |
| *lin-35* | HFW177 | *lin-35(n745)* | CGC |
| *xnp-1(EMS suppressor)* | HFW47 | *xnp-1(hcf3)* | This study |
| *toe-1* | HFW139 | *toe-1(syb6057);* | This study |
| *toe-1; xnp-1* | HFW153 | *xnp-1(tm678); toe-1(syb6057)* | This study |
| *mog-5* | HFW138 | *mog-5(syb5976)* | This study |
| *mog-5; xnp-1* | HFW144 | *mog-5(syb5976); xnp-1(tm678)* | This study |
| *hgap-2* | HFW158 | *hgap-2(syb7188)* | This study |
| *hgap-2; xnp-1* | HFW175 | *hgap-2(syb7188); xnp-1(tm678)* | This study |
| *gsox-1* | HFW159 | *F33H1.4(syb7245)* | This study |
| *gsox-1; xnp-1* | HFW176 | *F33H1.4(syb7245); xnp-1(tm678)* | This study |
