## supplementary table 2 for "ATRX safeguards cellular identity during *C. elegans* development"

**Supplementary Table Two**

| Name | Primers Used (5’ – 3’) | Primer Source |
| --- | --- | --- |
| *Cer8* | F: GCACGAATGT  GAGCTCTCCG  R: CCCATTGTC  GGGACTTTCG | This study |
| *Cer10* | F: TACCAACGAG  CCGAGTCTTC  R: TCTTCAGTTT  CCTCGCCTGT | Zeller *et al.*, 2016 |
| *tba-1* | F: TCAACACTGC  CATCGCCGCC  R: TCCAAGCGAGA  CCAGGCTTCAG | Zhang *et al.*, 2012 |
| *LINE2A* | F: CCTGCTAACTA  CCACTTCATACTG  R: CGGTACGAG  TGGGCTTC | This study |
| *Vingi-2* | F: CAACATGAAGC  AGCCG  R: GGTTCCCAA  AGTTAGCTTTG | This study |
| *RTE1* | F: CGTTTCTCCCG  GAAGAAATTCG  R: GGGTTTCGGTAC  ATTTCTGCTGT | This study |
| *CELE45* | F: TCCACCCAGTTT CTATTGAGAAGG  R: TGAACTGTTCTT  CAAGCGCTATG | This study |
| *CEMUDR* | F: AGGCCCATTC  CATAGCTTTT  R: TTTCTGGGAT  TTCATGCACA | Zeller *et al.*, 2016 |
| *Marince1* | F: CACAGTCACT  ACACCAGCG  R: GTGAGCTCCAT  TTCTGAATCTTCGG | This study |
| *LONGPAL2* | F: CTACGATTAAC  GATTTTTACTACC  R: CGGATTCAGT  TAAAAACCGC | This study |
| *pgl-1* | F: TGTTGTTGGA  GTCGCGAAG  R: TCCGCAATG  GCTCGTCTT | Updike and Strome, 2009 |

Zeller P, Padeken J, van Schendel R, Kalck V, Tijsterman M, Gasser SM. Histone H3K9 methylation is dispensable for Caenorhabditis elegans development but suppresses RNA:DNA hybrid-associated repeat instability. Nat Genet. 2016 Nov;48(11):1385-1395. doi: 10.1038/ng.3672. Epub 2016 Sep 26.

Zhang Y, Chen D, Smith MA, Zhang B, Pan X. Selection of reliable reference genes in Caenorhabditis elegans for analysis of nanotoxicity. PLoS One. 2012;7(3):e31849. doi: 10.1371/journal.pone.0031849. Epub 2012 Mar 15.

Updike DL, Strome S. A genomewide RNAi screen for genes that affect the stability, distribution and function of P granules in Caenorhabditis elegans. Genetics. 2009 Dec;183(4):1397-419. doi: 10.1534/genetics.109.110171. Epub 2009 Oct 5.
