## supplementary figure legends for "ATRX safeguards cellular identity during *C. elegans* development"

**Supplementary Figure One – Loss of XNP-1 reduces the synchrony of *C. elegans* development.**

Heatmaps displaying trend-corrected luminescence signal (arbitrary units) over time for wildtype and mutant animals that express luciferase from a ubiquitous and constitutive promoter (*eft-3*). Each row represents a single animal, with timepoint 0 representing the moment of hatching. Molts are defined by extended periods with darker signal, associated with lower luminescence, and individual animals in each heatmap are sorted based on time of entry into the first molt.

**Supplementary Figure Two – No change in the levels of 22G RNAs targeting upregulated repetitive elements in *xnp-1***

(A) Loss of XNP-1 leads to a modest upregulation of repetitive elements in both embryos and L1 larvae. Embryos and L1 animals were grown at 20°C and processed for polyA-RNA sequencing. The expression of the indicated transposons is displayed as a heatmap for each biological replicate. The adjacent graph displays the log_2_ fold change (FC) plotted with the data normalised to the *xnp-1(tm678)* strain in a stage-specific manner and to the protein-coding gene reads on the x-axis compared to the -log_2_ of the false discovery rate (FDR). Significantly expressed transposon families are marked in purple. (B) Small RNAs (sRNAs) were isolated from the same total RNA as the polyA-enriched samples in Figure 2A. The graphs display the relative fold change in 22G RNAs targeting the transposable elements whose mRNA expression increases in *xnp-1* in either embryos or L1 larvae (Figure 2A). The abundance of these 22G RNAs is unchanged in *xnp-1*, indicating that the change in transposable element mRNA levels is not driven by the RNAi (22G RNA) pathway.

**Supplementary Figure Three – Effect on XNP-1 on Cer10 is independent of H3K9me3**

(A) RT-qPCR of *Cer10* with *tba-1* used as the reference in wildtype and *xnp-1(tm678)* animals with *Cer10* expression normalised to wildtype. The mean and standard deviation is shown for three biological replicates. * = p ≤ 5 x 10^-2^, (two tailed T-test). The ability of *xnp-1* mutation to increase *Cer10* expression is independent of H3 K9me3 (SET-25).

**Supplementary Figure Four – *xnp-1* does not phenocopy *lin-35***

(A) Comparison of genes upregulated in *lin-35* in L1 larvae (Petrella *et al*., 2011) with genes upregulated in *xnp-1* (Figure 2A) shows little overlap. (B) *mes-4(RNAi)* in *lin-35* shows the expected partial rescue of larval arrest at 25°C. This acts as a positive control for RNAi knockdown in Figure 2F. Genes downregulated in xnp-1 embryos or L1s at 20C are not significantly enriched for particular gene ontology terms but are significantly associated with the intestine and rectal valve cell.

**Supplementary Figure Five – *NHR-23 binding profiles are not similar to GSOX-1***

(A) ChIP-seq data of NHR-23 binding (Kudron *et al*., 2024) analysed with deepTools indicating that in contrast to GSOX-1 (Figure 4E) it displays similar levels of binding at promoters of genes upregulated or downregulated in *xnp-1.*
